# Development and Validation Molecular Docking Analysis of Human serum albumin (HSA)

**DOI:** 10.1101/2021.07.09.451789

**Authors:** Ivan Vito Ferrari, Paolo Patrizio

## Abstract

**Background:** HAS (Human Serum Albumin) is a highly water-soluble globular plasma protein, with a relative molecular weight (g/mol) of 67 KDa, consisting of 585 amino acid residues. In this study, we have investigated the interaction of the Crystal structures complexed in human serum albumin at resolutions of 2.8 to 2.0: Camptothecin, 9-amino-camptothecin, Etoposide, Teniposide, Bicalutamide and Idarubicin, using a bioinformatic approach, estimated by Pyrx Virtual Screen Tool and AMDock (AMDock, Assisted Molecular Docking). We have analyzed a validated protocol, studying several parameters, as Binding Affinity, RMSD value, Ligand Efficiency, and Inhibition constant (Ki value).

**Methods:** Human Serum Albumin protein preparation was characterized with several programs, as Chimera, MGLTools 1.5.6, Swiss PDB Viewer Software to perform docking analysis by Autodock Vina estimated with Pyrx Software.

**Results:** In this work, we have found crystalized camptothecin, crystalized 9-amino-camptothecin and crystalized teniposide, gave excellent results for Binding Affinity, (kcal/mol), RMSD value (A°), inhibition constant Ki value (nM): -<u>Binding Affinity</u> of 9-amino-camptothecin (ca.−10 kcal/mol), camptothecin (−9 kcal/mol) and teniposide (−11 kcal/mol, -<u>RMSD Value</u> of 9 -amino-camptothecin (ca.1.8 Å), camptothecin (ca.2.2 Å) and teniposide (ca. 3.6 Å), - <u>Ki Value</u>: 9 -amino-camptothecin (ca 59 nM), camptothecin (ca 183 nM) and teniposide (ca 9 nM), -<u>Ligand efficiency</u>: of 9 -amino-camptothecin(ca −0.35 kcal/mol), camptothecin (ca −0.34 kcal/mol) and teniposide (ca −0.24 kcal/mol

**Conclusions:** We explored the best three crystallized ligand in Human Serum Albumin. Moreover, we observe a complete overlap, during the re-docking analysis phase, estimated by chimera Software. Therefore we have concluded that ID PDB Crystal 4L8U human serum albumin-Crystallised 9 -amino Camptothecin; ID PDB Crystal 4L9K human serum albumin-Crystallised Camptothecin and ID PDB Crystal 4L9Q human serum albumin-crystallized teniposide be used as a possible as a reference template protein to be compared with the target protein, by Docking molecular analysis.

## 1 Introduction

HAS (Human Serum Albumin) is a highly water-soluble globular monomeric plasma protein, with a relative molecular weight (g/mol) of 67 KDa, consisting of 585 amino acid residues. It is the most abundant protein in human blood, acting as a natural drug carrier and it has many vital biological roles. Moreover, HAS plays a fundamental role in the transport of drugs, metabolites, and endogenous ligands. Albumin made by the liver is a family of globular proteins, the most common of which is the serum albumins The gene for albumin is located on chromosome 4 in locus 4q13.3. Serum albumin possesses a unique capability to bind, covalently or reversibly, a great number of various endogenous and exogenous compounds. In fact, Albumin transports hormones, fatty acids, and drugs. The reference range for albumin concentrations in serum is approximately 35-50 g/L (3,5-5,0 g/Dl). It has a serum half-life of approximately 21 days. Albumin is synthesized in the liver as preproalbumin, which has an N-terminal peptide that is removed before the nascent protein is released from the rough endoplasmic reticulum. The product, proalbumin, is in turn cleaved in the Golgi apparatus to produce the secreted albumin. Serum albumin interaction with drug molecule has been an essential parameter to understand the pharmacokinetic properties of a drug such as a half-life, solubility, distribution, and metabolism ^1–15^. HSA consists of a single polypeptide chain of 585 amino acids but with three homologous domains: domain I (residue 1–195), II (residue 196–383) and III (residue 384–585) Each domain possesses two subdomains A and B formed by six and four α-helices, respectively. The hydrophobic pocket of subdomain IIA and IIIA, popularly known as Sudlow’s site I and II, respectively accommodate the aromatic and heterocyclic compounds ^16–26^. Here we investigate, through a study in Silico Bioinformatic approach, the interaction of the crystal structures of six oncology agents were determined in complex with human serum albumin at resolutions of 2.8 to 2.0 Å: camptothecin, 9-amino-camptothecin, etoposide, teniposide, bicalutamide, and idarubicin, using a bioinformatic approach, by Molecular Docking Analysis by AutoDock Vina Tool, estimated Pyrx Virtual Screen Tool and AMDock (AMDock (Assisted Molecular Docking) Software, taking as a reference model a previous paper of Zhong-min Wang, et, al., (2013) ^11^. The main idea of this paper is to focus on the new development and validation method, in the Molecular Docking analysis of the crystal structures of six oncology agents. They were have investigated studying: Binding Affinity (kcal/mol), RMSD value, Ligand Efficiency (kcal/mol), and inhibition constant (Ki value). Generally speaking, computational chemistry and chemoinformatics play a key role in early phase drug research, by identifying the most promising candidates for experimental investigations. Two major strategies have been employed for virtual screening: pharmacophore modeling and molecular docking. Virtual screening (VS) is a computer-aided method to simulate High-throughput screening (HTS) that can save time and costs in the drug development process, also reducing the failure rate by prioritizing compounds for further experimental investigation. HTS is a method to identify new leads for drug discovery that allows a large chemical library to be screened in vitro against a specific drug target, cell, or organism. Docking is a molecular modeling method that allows compounds to be screened in silico before being tested experimentally. Ligand docking is a widely used approach in virtual screening ^26—29^. In figure 1–3 we report Human serum albumin (HSA) structure as cartoon representation and several ID PDB protein crystal structures of six oncology agents were determined in complex with human serum albumin: a) ID PDB 2.40 Å Resolution: Crystal 4LA0 human serum albumin-Crystallised Bicalutamide b) ID PDB 2.01 Å Resolution: Crystal 4L8U human serum albumin-Crystallised 9 amino Camptothecin c) ID PDB 2.40 Å: Crystal 4L9K human serum albumin-Crystallized Camptothecin d) ID PDB 2.70 Å Resolution: Crystal 4LB9 human serum albumin-Crystallised etoposide e) ID PDB 2.80 Å Resolution: Crystal 4LB2 human serum albumin-Crystallised idarubicin f) ID PDB 2.70 Å Resolution: Crystal 4L9Q human serum albumin-Crystallised teniposide ^11^.

**Fig 1.**
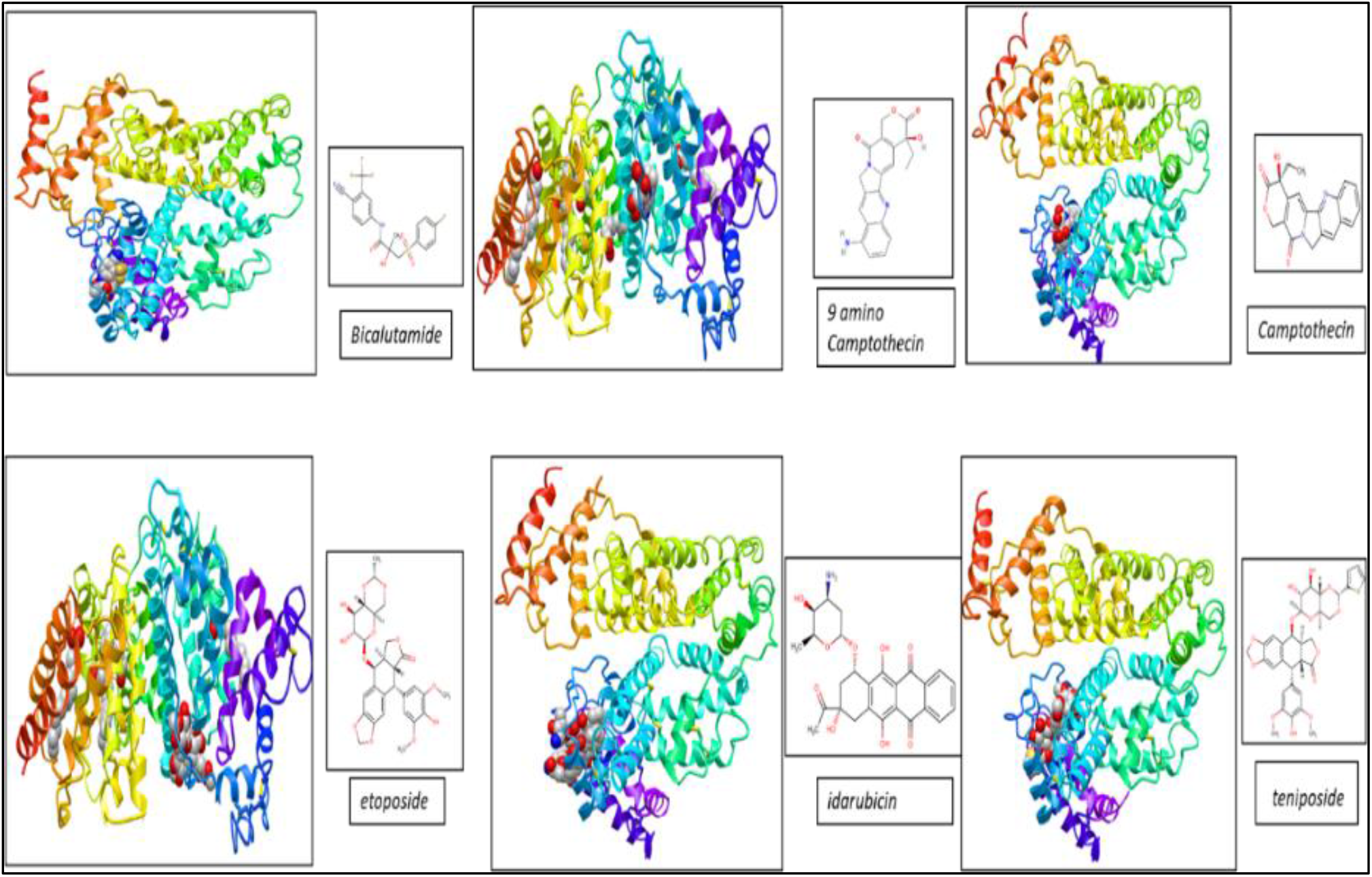
ID PDB 2.40 Å Resolution : Crystal 4LA0 human serum albumin-Crystallised Bicalutamide b) ID PDB 2.01 Å Resolution: Crystal 4L8U human serum albumin-Crystallised 9 amino Camptothecin c) ID PDB 2.40 Å Resolution: Crystal 4L9K human serum albumin-Crystallised Camptothecin d) ID PDB 2.70 Å Resolution: Crystal 4LB9 human serum albumin-Crystallised etoposide e) ID PDB 2.80 Å Resolution: Crystal 4LB2 human serum albumin-Crystallised idarubicin f) ID PDB 2.70 Å: Crystal 4L9Q human serum albumin-Crystallised teniposide ^11;^ reproduced by chimera software

**Fig 2.**
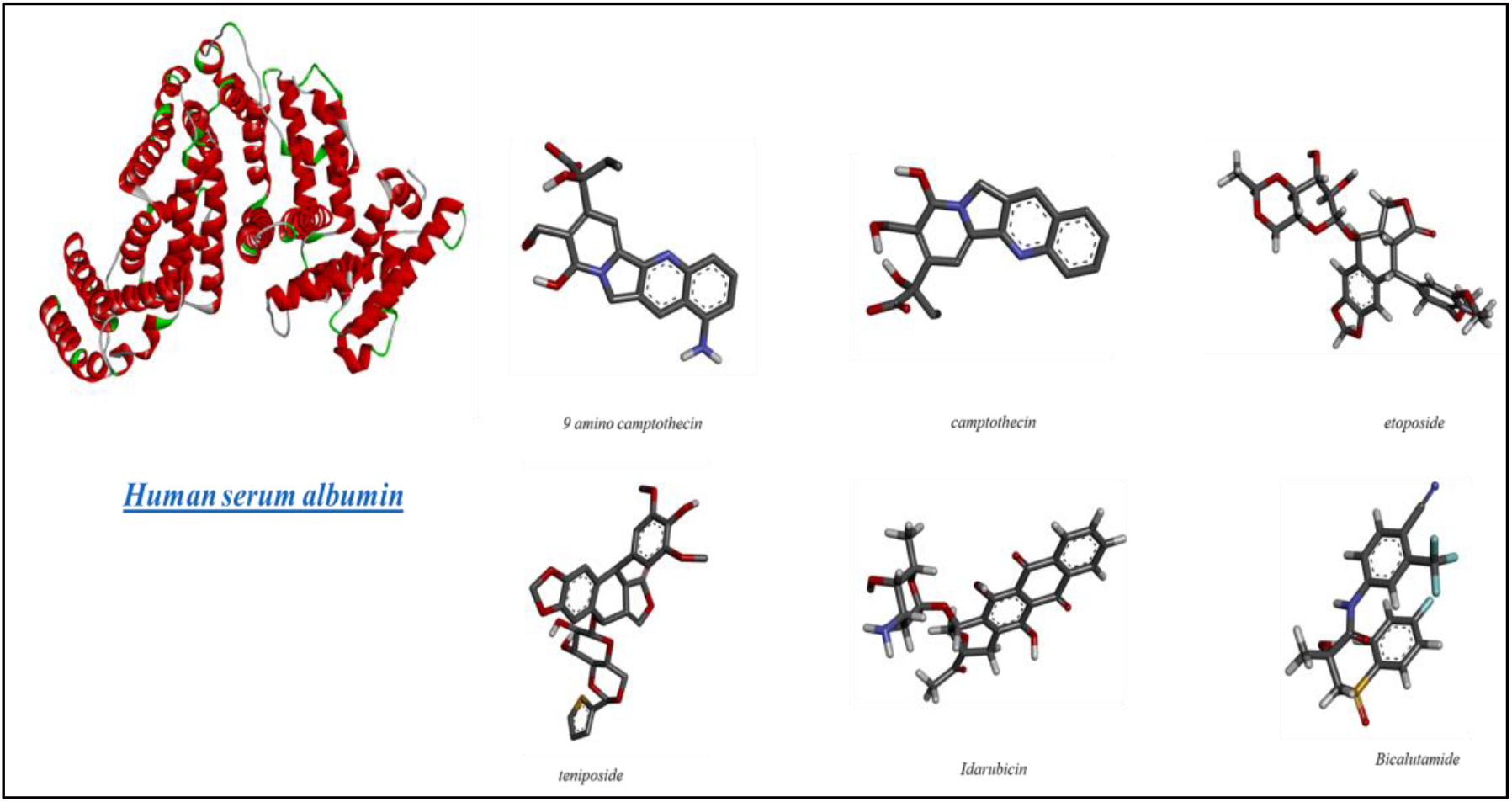
On the right Chemical Structure of six oncology agents: 9 amino Camptothecin; Camptothecin; etoposide; teniposide idarubicin and bicalutamide and on the left Human serum albumin structure as cartoon reproduced by Discovery Studio Biovia software

**Fig 3.**
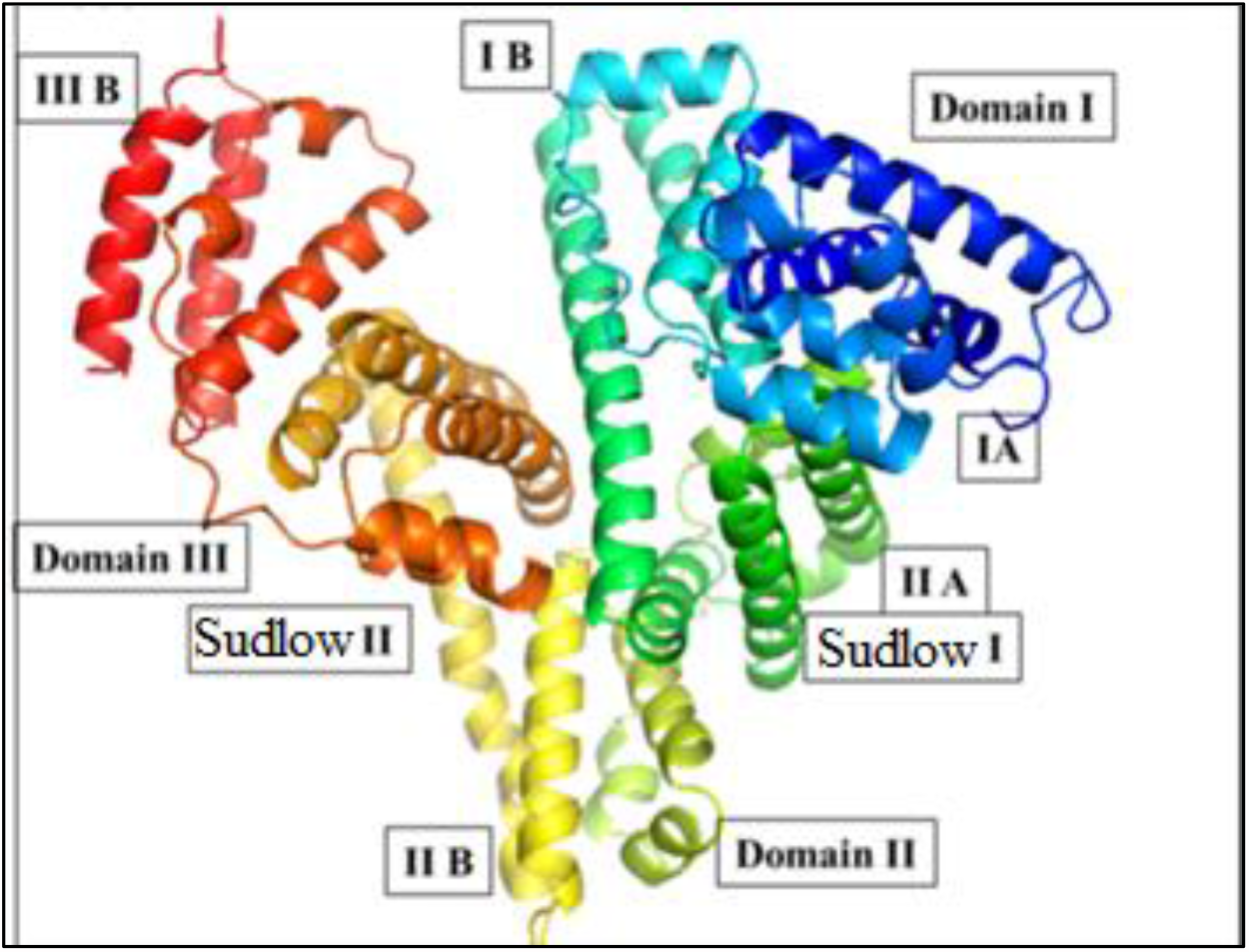
The structure of Human serum albumin (HSA) representing domains, subdomains and Sudlow’s binding sites I and II. ^24–25^ Reproduced by Pymol software

## 2. Materials and methods

### 2.1. Human serum albumin (HSA)

We have reported Human serum albumin (HSA) structure as cartoon representation, in PDB format. It has been created by downloading from the protein data bank (https://www.rcsb.org). HSA consists of a single polypeptide chain of 585 amino acids but with three homologous domains: domain I (residue 1–195), II (residue 196–383), and III (residue 384–585) Each domain possesses two subdomains A and B formed by six and four α-helices, respectively. The hydrophobic pocket of subdomain IIA and IIIA, popularly known as Sudlow’s site I and II, respectively accommodate the aromatic and heterocyclic compounds. Structurally, albumin is a 585 amino acid protein. 11These three gene domains 69 are homologous (I, II, III). Each domain in turn is comprised of two subdomains, denoted A and B 25-26 Meloun B, et all., (1975) has investigated the amino acid analysis of human serum albumin, showing that the molecule of this protein contains approximately 575 amino acid residues, of their number 6 methionines ^32^.

#### 2.3.1. Protein Preparation

The three-dimensional crystal structure of Human serum albumin (HAS) ID PDB 4LA0 human serum albumin-Crystallised Bicalutamide; ID PDB 4L8U human serum albumin-Crystallised 9 amino Camptothecin; ID PDB 4L9K human serum albumin-Crystallised Camptothecin; ID PDB 4LB9 human serum albumin-Crystallised etoposide; ID PDB 4LB2 human serum albumin-Crystallised idarubicin and ID PDB; 4L9Q human serum albumin-Crystallised teniposide in PBD format from the protein data bank was accomplished.

##### Workflow protein preparation steps

ID PDB 4LA0, ID PDB 4L8U, ID PDB 4L9K, ID PDB 4LB9, ID PDB 4LB2 and ID PDB 4L9Q were downloaded from the protein data bank (https://www.rcsb.org) and saved in pdb format. The following steps: a) delete all co-crystallized ligands and waters in the protein by Chimera Software b) Add Kollmann charges and add only Polar Hydrogen in the protein, by MGLTools 1.5.6/ (c) replaced missing residues and energy minimization step by Swiss PDB Viewer software. Final step, the protein has been converted into pdbqt format ready for docking process by Pyrx Software. (See below figure 4)

**Fig 4.**
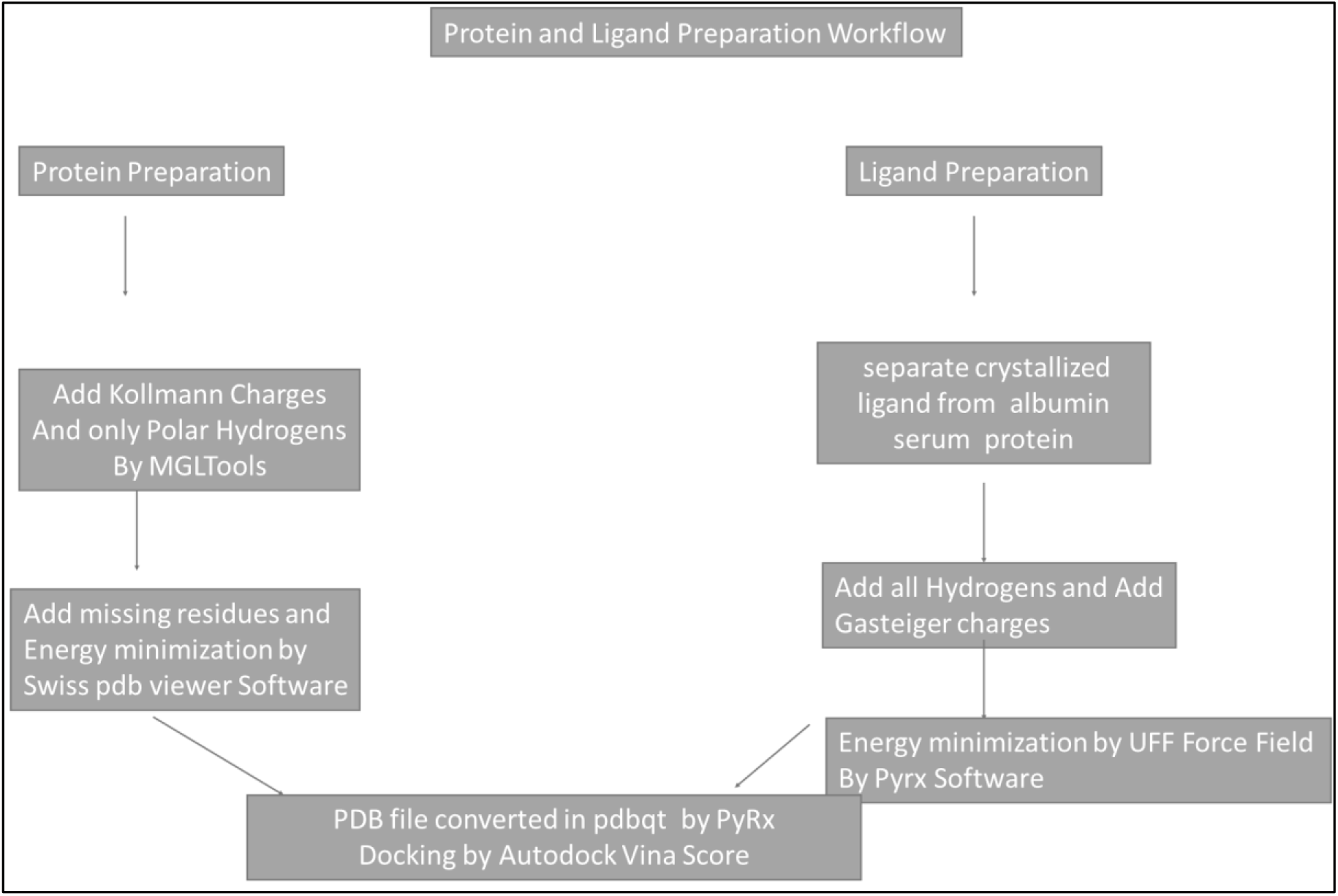
Workflow Protein and Ligand preparation steps prior to performe molecular docking analysis by AutoDock vina estimated by Pyrx and AMDock

#### 2.3.2 Ligand Preparation

##### Workflow Ligand preparation steps

the six oncology agents (Crystallised 9 amino Camptothecin; Camptothecin; etoposide; teniposide, idarubicin, and bicalutamide a) were separated individually from the protein b) add all hydrogen and add charge Gasteiger by MGLTools 1.5.6 c) Energy minimization step by 4 force field (uff) method, by Pyrx Virtual Screening Tool d) drug structure files have been converted into pdbqt format, Pyrx Software, and AMDock Software. (See below figure 4)

### 2.4. Pyrx Virtual Screening Tool

PyRx is a Virtual Screening software for Computational Drug Discovery. It includes chemical spreadsheet-like functionality and powerful visualization engine that are essential for structure-based drug design. In this paper we performe Molecular docking analysis, in order to estimate the Affinity (kcal/mol) ^35–36^. Binding affinity (Kcal/mol) of ID PDB: 4L8U human serum albumin (HSA) - complexed with Crystallised 9 amino camptothecin was calculated in: [central position and size of the grid docking box: center x=29.23;y=5.84;z=31.39/ size x,y,z= 25/25/25 Å]. ID PDB: 4L9K human serum albumin (HSA) - complexed with Crystallised camptothecin: was calculated in: [central position and size of the grid docking box: x=−9.85; y=−6.88; z=9.79/ size x,y,z= 25/25/25 Å]. ID PDB: 4L9Q human serum albumin (HSA) - complexed with Crystallised teniposide was calculated in: [central position and size of the grid docking box: x=−13.107; y=−11.28;z=9.95/ size x,y,z= 25/25/25 Å]. ID PDB: 4LA0 human serum albumin (HSA) - complexed with Crystallised bicalutamide was calculated in: [central position and size of the grid docking box: x=−16.45; y=−12.02; z=8.18/ size x,y,z= 25/25/25 Å]. ID PDB: 4LB2 human serum albumin (HSA) - complexed with Crystallised idarubicin was calculated in: [central position and size of the grid docking box: x=−14.40; y=−14.24;z=9.80/ size x,y,z= 25/25/25 Å]. ID PDB: 4LB9 human serum albumin (HSA) - complexed with Crystallised etoposide was calculated in: central position and size of the grid docking box: x=33.40; y=−2.79;z=16.69/ size x,y,z= 25/25/25 Å].

### 2.5. AMDock (AMDock (Assisted Molecular Docking)

AMDock (Assisted Molecular Docking) is a user-friendly graphical tool to assist in the docking of protein-ligand complexes using AutoDock Vina and AutoDock4, including the option of using the Autodock4Zn force field for metalloproteins. AMDock integrates several external programs (Open Babel, PDB2PQR, Auto Ligand, ADT scripts).^48^ AMDock was used to calculate three important parameters of six oncology agents: (9 amino Camptothecin; Camptothecin; Etoposide; Teniposide, Idarubicin, and Bicalutamide) in Human serum albumin: Binding Affinity (kcal/mol), Ligand Efficiency and inhibition constant (Ki value) Ki is a measurement of the binding energy per atom of a ligand to its binding partners, such as a receptor or enzyme. Binding affinity is the strength of the binding interaction between a single biomolecule (e.g. protein or DNA) to its ligand/binding partner (e.g. drug or inhibitor). Ligand efficiency (LE) is used in drug discovery research programs to assist in narrowing the focus to lead compounds with optimal combinations of physicochemical properties and pharmacological properties. Binding affinity is typically measured and reported by the equilibrium dissociation constant (KD), which is used to evaluate and rank order strengths of biomolecular interactions.

The smaller the KD value, the greater the binding affinity of the ligand for its target. The inhibitory constant (Ki) is the concentration of the inhibitor that is required to decrease the maximal rate of the reaction by half ^48^.

### 2.6. AutoDock 4 and AutoDock Vina

AutoDock Vina is an open-source program for doing molecular docking. It was designed and implemented by Dr. Oleg Trott in the Molecular Graphics Lab at The Scripps Research Institute. AutoDock is a suite of automated docking tools. Current distributions of AutoDock consist of two generations of software: AutoDock 4 and AutoDock Vina. AutoDock Vina is a new generation of docking software from the Molecular Graphics Lab. It is very fast, provides high-quality predictions of ligand conformations, and good correlations between predicted inhibition constants and experimental ones. In AutoDock 4 the scoring function is based on the AMBER force field and in AutoDock Vina it is a hybrid scoring function (empirical + knowledge-based) based on the X-Score function with some different parameters which is not published at the moment. AutoDock Vina is much faster and more accurate (depending on the system). It calculates the grid charges internally and setting up the docking is much easier. Adding charges depends on your method and not the program used. More accurate charge calculations can lead to better docking results but they are more computationally expensive. AutoDock Vina ignores user-supplied charges ^37–43^.

### 2.7. BIOVIA Discovery Studio Visualizer Software

BIOVIA Discovery Studio is a comprehensive suite of validated science applications built on BIOVIA Pipeline Pilot. This software was used to create a 2D diagram of Ligand Binding Site Atoms of six oncology agents: 9 amino Camptothecin; Camptothecin; Etoposide; Teniposide, Idarubicin, and Bicalutamide in Human serum albumin ^44–45^.

### 2.8. UCSF Chimera: An Extensible Molecular Modeling System

UCSF Chimera is a program for the interactive visualization and analysis of molecular structures and related data, including density maps, trajectories, and sequence alignments. This Software was used in order to re-investigation Binding affinity and validation method docking Score by AutoDock Vina. This Software was used in order to re-investigated Binding affinity and to calculate RMSD value of six oncology agents: 9 amino Camptothecin; Camptothecin; Etoposide; Teniposide, Idarubicin and Bicalutamide in Human serum albumin ^46–47^.

### 3.0. Binding affinity

Molecular docking is a key tool in structural molecular biology. Protein−ligand binding is essential to almost all biological processes. Binding affinity is the strength of the binding interaction between a single biomolecule (e.g. protein or DNA) to its ligand/binding partner (e.g. drug or inhibitor). Binding affinity is typically measured and reported by the equilibrium dissociation constant (KD), which is used to evaluate and rank order strengths of biomolecular interactions. The smaller the KD value, the greater the binding affinity of the ligand for its target. Low-affinity binding (high Ki level) implies that a relatively high concentration of a ligand is required before the binding site is maximally occupied and the maximum physiological response to the ligand is achieved^49–53^.

The AutoDock semi-empirical force field includes intramolecular terms. Water molecules are not modelled explicitly though, but pair-wise atomic terms are used to estimate the water contribution (dispersion/repulsion, hydrogen bonding, electrostatics, and desolvation), where weights are added for calibration (based on experimental data). The evaluation step in a nutshell: firstly, calculate the energy of ligand and protein in the unbound state. Secondly, calculate the energy of the protein-ligand complex ^49–53^.

### 2. 1. RMSD (Root-mean-square deviation)

The study of RMSD for small organic molecules (commonly called ligands when they’re binding to macromolecules, such as proteins, is studied) is common in the context of docking as well as in other methods to study the configuration of ligands when bound to macromolecules. An RMSD value is expressed in length units. The most commonly used unit in structural biology is the Angstrom (Å) which is equal to 10−10 m ^54^.

## 3. Results and discussion

### 3.1. Validation of Molecular Docking

We performed a docking investigation of six oncology crystal structures determined in complex with human serum albumin at resolutions of 2.8 to 2.0 Å: Camptothecin, 9-amino-Camptothecin, Etoposide, Teniposide, Bicalutamide, and Idarubicin. In detail, we used ID PDB 2.40 Å Resolution: Crystal 4LA0 human serum albumin-Crystallised Bicalutamide b)

ID PDB 2.01 Å Resolution: Crystal 4L8U human serum albumin-Crystallised 9 amino Camptothecin c) ID PDB 2.40 Å Resolution: Crystal 4L9K human serum albumin-Crystallised Camptothecin d) ID PDB 2.70 Å Resolution: Crystal 4LB9 human serum albumin-Crystallised etoposide e) ID PDB 2.80 Å Resolution: Crystal 4LB2 human serum albumin-Crystallised idarubicin f) ID PDB 2.70 Å Resolution 4L9Q human serum albumin-Crystallised teniposide. It should be borne in mind that there are many methods reported for validating docking programs and scoring functions. Programs that can return poses below a preselected Root Mean Square Deviation (RMSD) value from the known conformation (usually 1.5 or 2 Å depending on ligand size) are considered to have performed successfully is RMSD (Root-mean-square deviation). The root-mean-square deviation or root-mean-square error is a frequently used measure of the differences between values predicted by a model or an estimator and the values observed. The RMSD represents the square root of the second sample moment of the differences between predicted values and observed values or the quadratic mean of these differences.47-4 Generally speaking a good method of Validation of Molecular Docking is to separate the crystallized ligand from its protein and repeat docking analysis in the same area, where the crystallized ligand was located before. In this way, we can check quickly, whether the crystallized ligand and the same docked ligand are overlapping or not, through the value of parameter RMSD. On the whole, 2 Å RMSD for a pose is considered a docking success. In our case, we report in fig. 7–12 crystal structures determined in complex with human serum albumin with their redocking analysis.

**Fig. 5.**
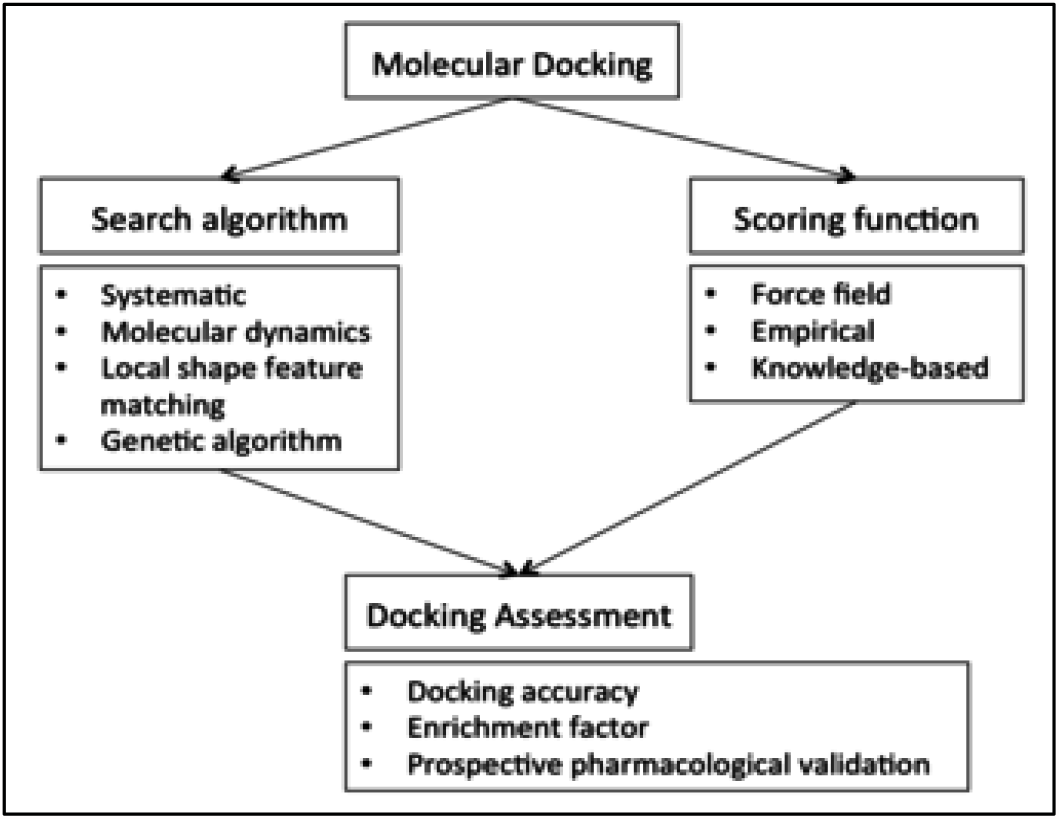
Docking flow-chart overview

**Fig.6.**
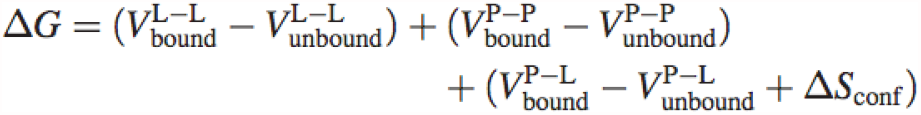
Where P refers to the protein, L refers to the ligand, V are the pair-wise evaluations mentioned above, and ΔS∼conf∼ denotes the loss of conformational entropy upon binding ^45–46^

**Fig 7.**
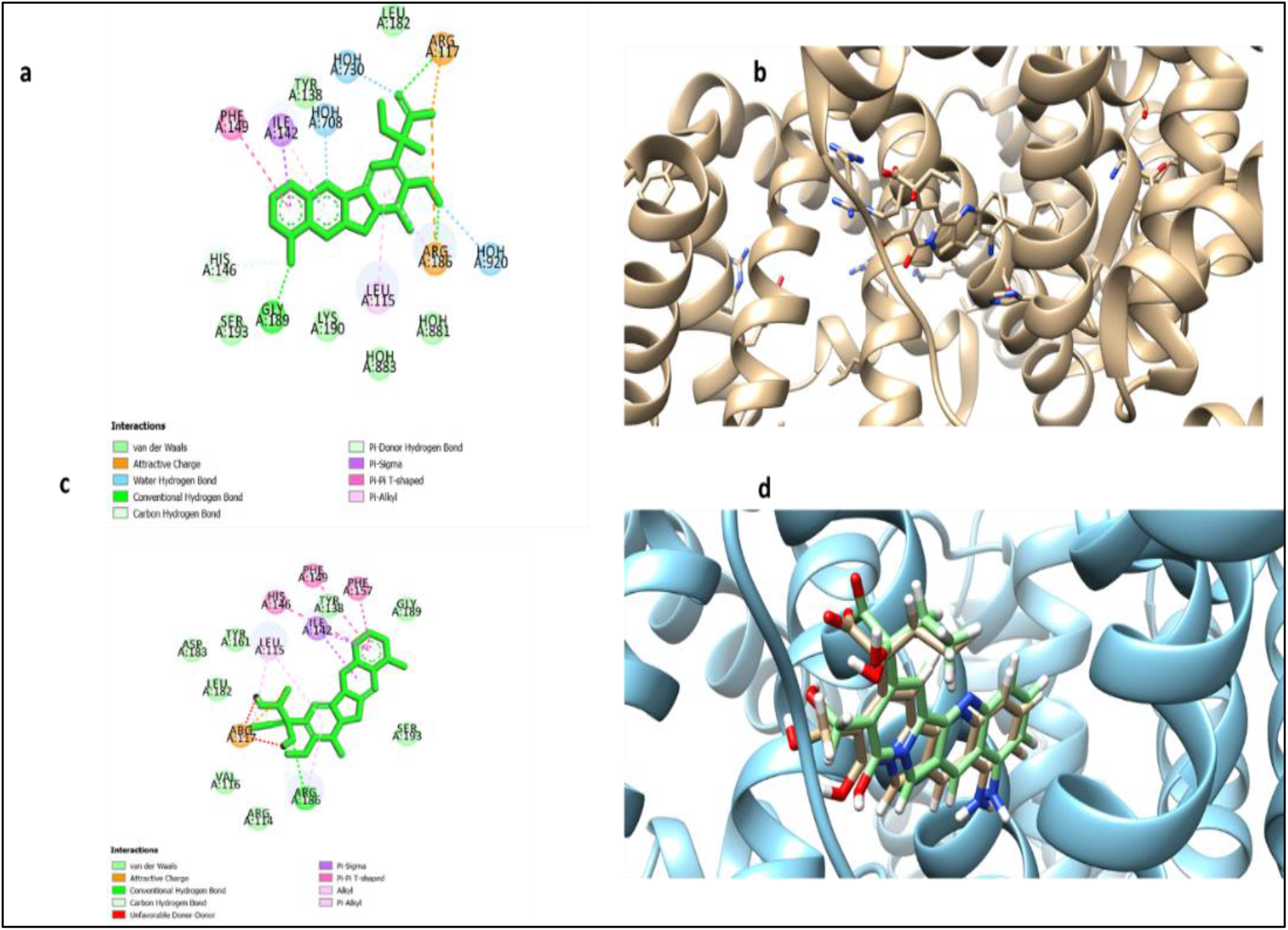
ID PDB: 4L8U human serum albumin (HSA) - complexed with Crystallised 9 amino camptothecin: a) 2D Diagram Ligand Binding site atoms using Discovery Studio Biovia b) protein-ligand complex using UCSF Chimera Software c) 2D Diagram Ligand Binding site atoms Re-Docked 9 amino camptothecin using Discovery Studio Biovia d) Comparison yellow Crystallised 9 amino camptothecin and green colour Re-Docked 9 amino camptothecin using UCSF Chimera Software [central position and size of the grid docking box: x=29.23;y=5.84;z=31.39/ size x,y,z= 25/25/25 Å]

**Fig 8.**
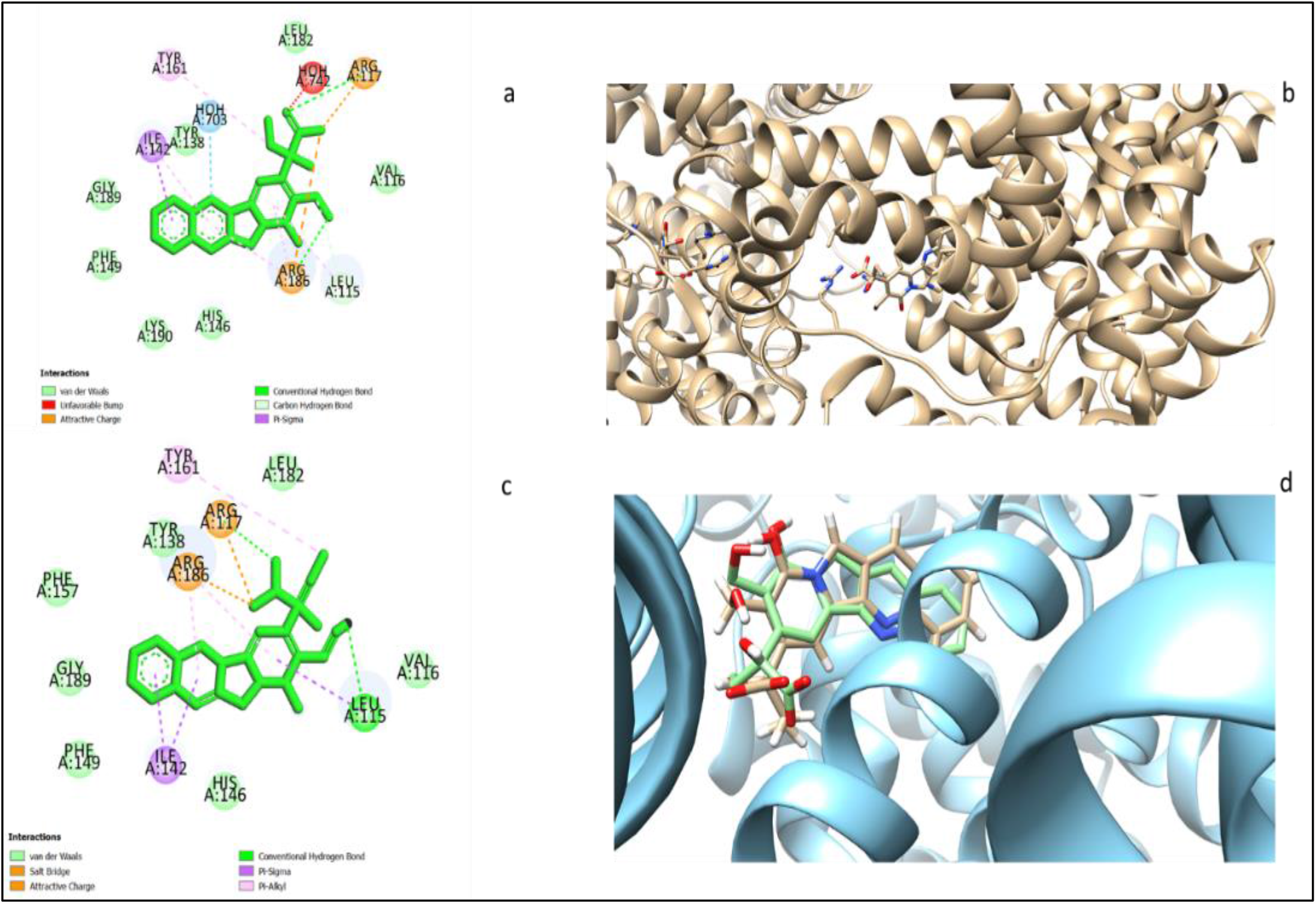
ID PDB: 4L9K human serum albumin (HSA) - complexed with Crystallised camptothecin: a) 2D Diagram Ligand Binding site atoms using Discovery Studio Biovia b) protein-ligand complex using UCSF Chimera Software c) 2D Diagram Ligand Binding site atoms Re-Docked camptothecin using Discovery Studio Biovia d) Comparison yellow Crystallised camptothecin and green colour Re-Docked camptothecin using UCSF Chimera Software [central position and size of the grid docking box: x=−9.85;y=−6.88;z=9.79/ size x,y,z= 25/25/25 Å]

**Fig 9.**
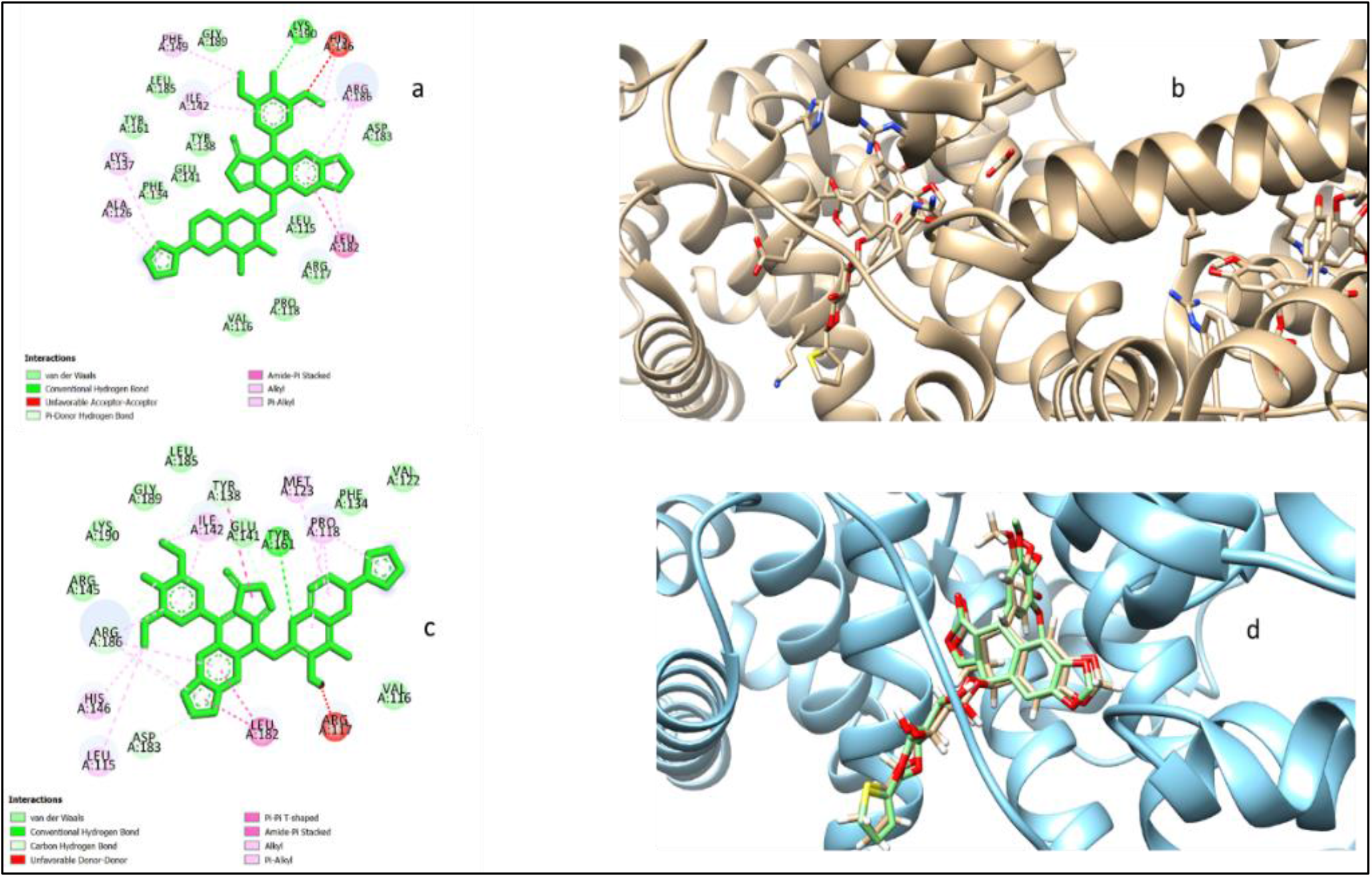
ID PDB: 4L9Q human serum albumin (HSA) - complexed with Crystallised teniposide a) 2D Diagram Ligand Binding site atoms using Discovery Studio Biovia b) protein-ligand complex using UCSF Chimera Software c) 2D Diagram Ligand Binding site atoms Re-Docked teniposide using Discovery Studio Biovia d) Comparison yellow Crystallised teniposide and green colour Re-Docked teniposide using UCSF Chimera Software[central position and size of the grid docking box: x=−13.107;y=−11.28;z=9.95/ size x,y,z= 25/25/25 Å]

**Fig 10.**
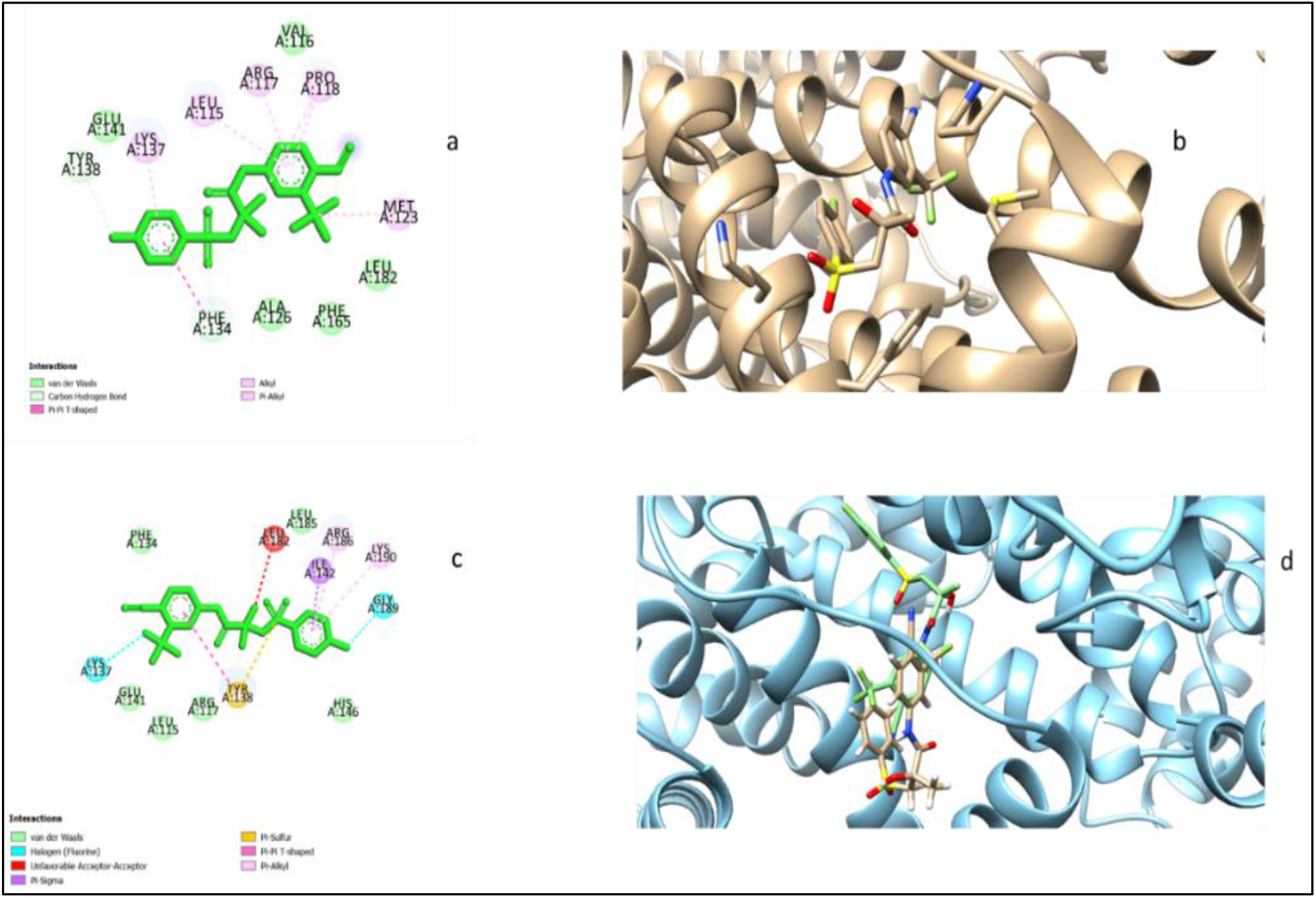
ID PDB: 4LA0 human serum albumin (HSA) - complexed with Crystallised bicalutamide a) 2D Diagram Ligand Binding site atoms using Discovery Studio Biovia b) protein-ligand complex using UCSF Chimera Software c) 2D Diagram Ligand Binding site atoms Re-Docked bicalutamide using Discovery Studio Biovia d) Comparison yellow Crystallised bicalutamide and green colour Re-Docked bicalutamide using UCSF Chimera Software[central position and size of the grid docking box: x=−16.45;y=−12.02;z=8.18/ size x,y,z= 25/25/25 Å]

**Fig 11.**
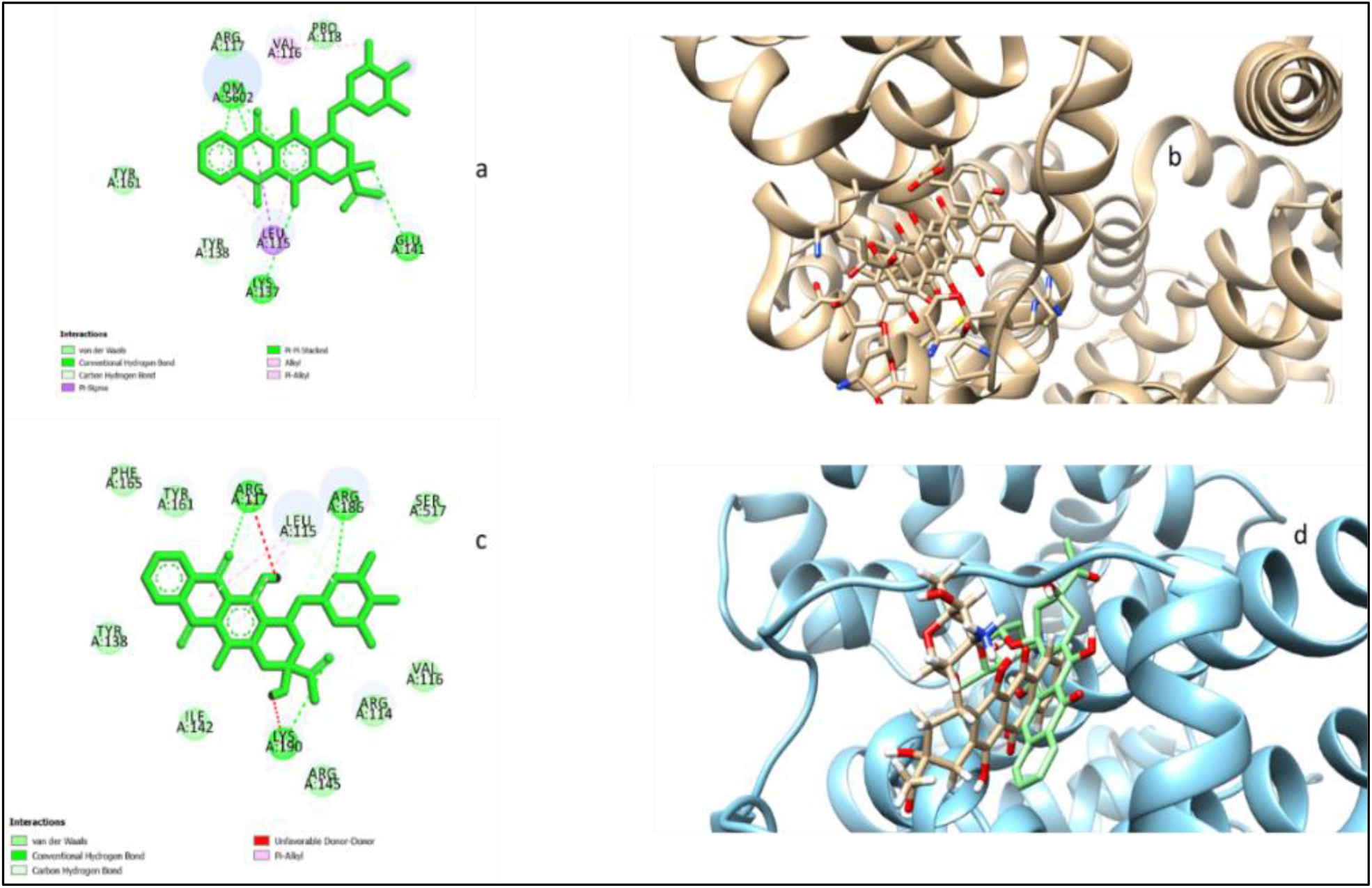
ID PDB: 4LB2 human serum albumin (HSA) - complexed with Crystallised idarubicin a) 2D Diagram Ligand Binding site atoms using Discovery Studio Biovia b) protein-ligand complex using UCSF Chimera Software c) 2D Diagram Ligand Binding site atoms Re-Docked idarubicin using Discovery Studio Biovia d) Comparison yellow Crystallised idarubicin and green colour Re-Docked idarubicin using UCSF Chimera Software[central position and size of the grid docking box: x=−14.40;y=−14.24;z=9.80/ size x,y,z= 25/25/25 Å]

**Fig 12.**
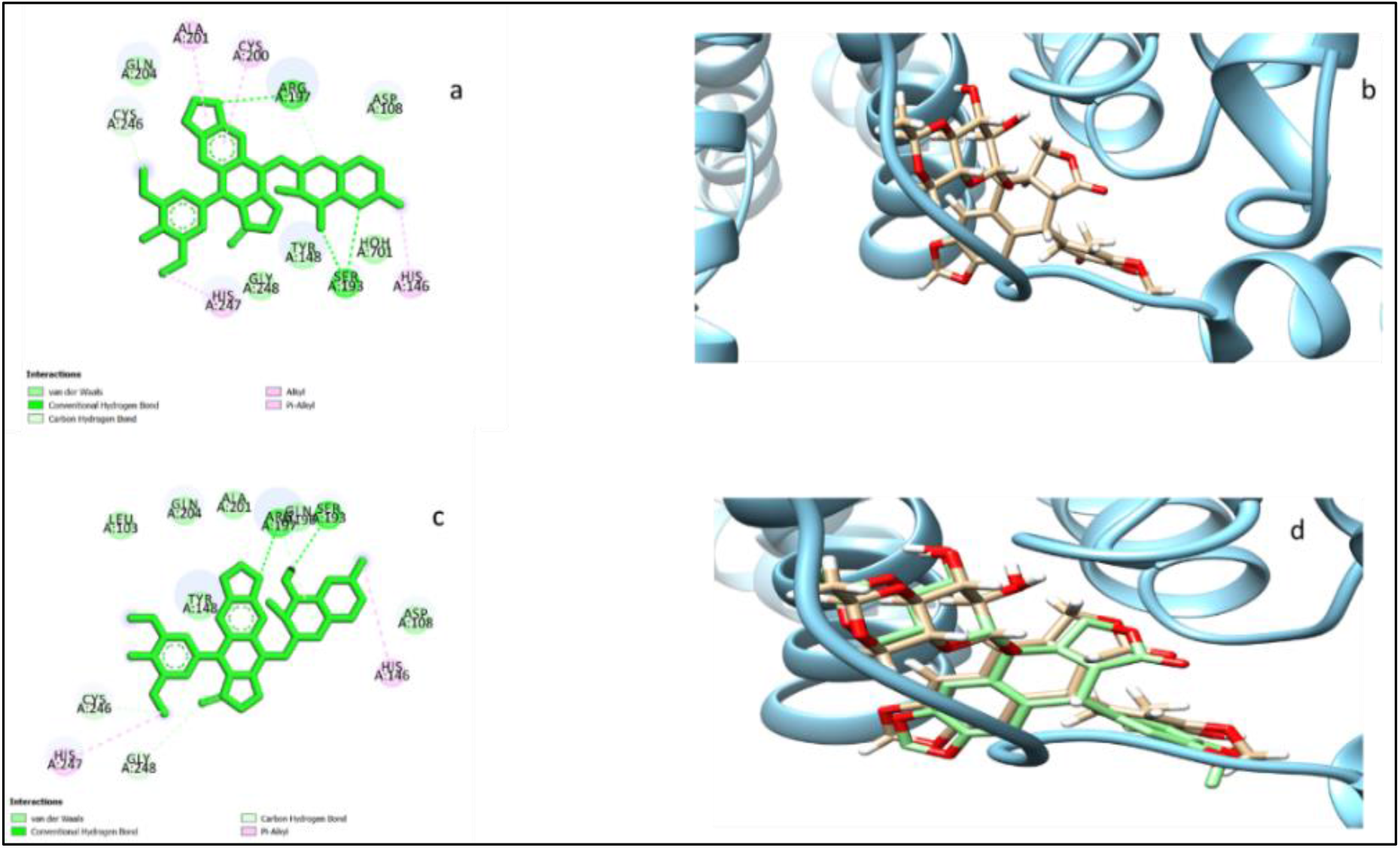
ID PDB: 4LB9 human serum albumin (HSA) - complexed with Crystallised etoposide a) 2D Diagram Ligand Binding site atoms using Discovery Studio Biovia b) protein-ligand complex using UCSF Chimera Software c) 2D Diagram Ligand Binding site atoms Re-Docked etoposide using Discovery Studio Biovia d) Comparison yellow Crystallised etoposide and green colour Re-Docked etoposide using UCSF Chimera Software[central position and size of the grid docking box: x=33.40;y=−2.79;z=16.69/ size x,y,z= 25/25/25 Å]

Moreover in fig. 7–12 we showed 2D Diagram Ligand Binding Site Atoms of six drugs: 9 amino Camptothecin; Camptothecin; Etoposide; Teniposide, Idarubicin, and bicalutamide, respectively, using Discovery Studio Biova Software.

In a particular way, we focused on residue interacting with H-bonding, Hydrophobic interactions, and van der Waals in human serum albumin. The main residues in ligand binding pocket site atoms involved are: Lys ^190^; Leu ^185^; Arg ^186^; His ^146^; Ile ^142^; Tyr ^161^; Tyr ^138^; Lys ^137^; Phe ^134^; Met ^123^, Pro ^118^; Arg ^117^; Val ^116^; Glu ^141^, Leu ^182^; Asp ^183 11^.

### 3.2. Molecular Docking of camptothecin, 9-amino-camptothecin, etoposide, teniposide, bicalutamide and idarubicin in HAS protein

In tab. 1–3 we have resumed the main results of six oncology agents investigated in human serum albumin. Binding Affinity (kcal/mol), was calculated by Pyrx Software, Chimera, and AMDock software. RMSD value (Root-mean-square deviation), was calculated by Chimera Software. Ligand Efficiency (kcal/mol) and inhibition constant (Ki value) were calculated by AMDock Software ((Assisted Molecular Docking). From the final results, we found that only three of the s crystallized six drugs in the plasma protein gave excellent results for Binding Affinity, RMSD value inhibition constant Ki value. They are camptothecin, 9-amino-camptothecin, and teniposide. In tab it was reported main results of these drugs:

**Tab 1.**
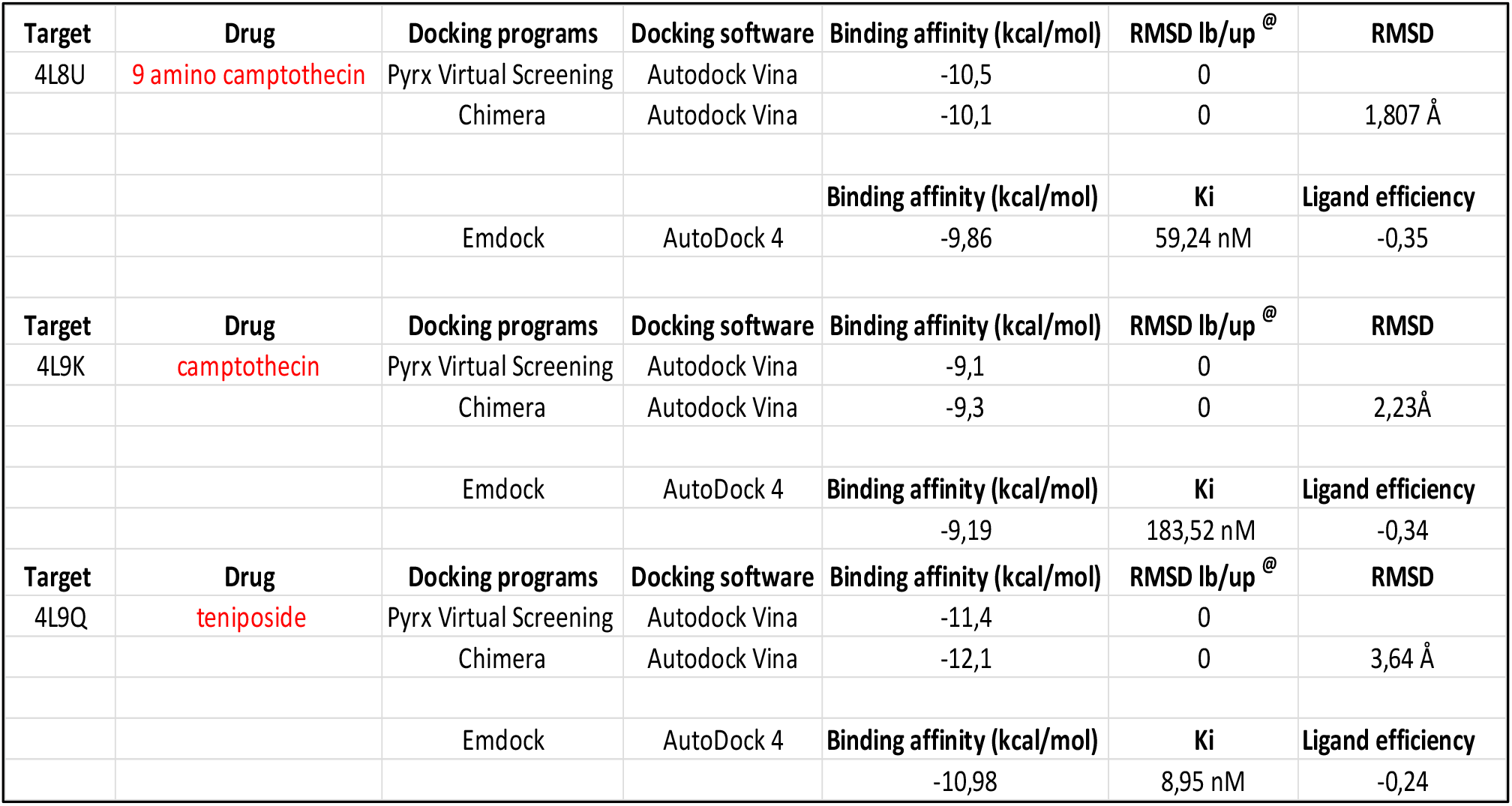
Catherization Docking analysis ID PDB:4L8U human serum albumin-Crystallised 9 amino Camptothecin; ID PDB: l 4L9K human serum albumin-Crystallised Camptothecin and ID PDB: 4L9Q human serum albumin-Crystallised teniposide

**Tab 2.** Docking of ID PDB: Crystal 4LA0 human serum albumin-Crystallised Bicalutamide b) ID PDB: Crystal 4L8U human serum albumin-Crystallised 9 amino Camptothecin c) ID PDB: Crystal 4L9K human serum albumin-Crystallised Camptothecin d) ID PDB: Crystal 4LB9 human serum albumin-Crystallised etoposide e) ID PDB: Crystal 4LB2 human serum albumin-Crystallised idarubicin f) ID PDB: Crystal 4L9Q human serum albumin-Crystallised teniposide

| Target | Drug | Docking programs | Docking software | Binding affinity (kcal/mol) | RMSD lb/up<sup>@</sup> | RMSD |
| --- | --- | --- | --- | --- | --- | --- |
| 4L8U | 9 amino camptothecin | Pyrx Virtual Screening | Autodock Vina | -10,5 | 0 |  |
|  |  | Chimera | Autodock Vina | -10,1 | 0 | 1,807 Å |
|  |  | Emdock | AutoDock 4 | Binding affinity (kcal/mol)<br>-9,86 | Ki<br>59,24 nM | Ligand efficiency<br>-0,35 |
| Target | Drug | Docking programs | Docking software | Binding affinity (kcal/mol) | RMSD lb/up<sup>@</sup> | RMSD |
| 4L9K | camptothecin | Pyrx Virtual Screening | Autodock Vina | -9,1 | 0 |  |
|  |  | Chimera | Autodock Vina | -9,3 | 0 | 2,23 Å |
|  |  | Emdock | AutoDock 4 | Binding affinity (kcal/mol)<br>-9,19 | Ki<br>183,52 nM | Ligand efficiency<br>-0,34 |
| Target | Drug | Docking programs | Docking software | Binding affinity (kcal/mol) | RMSD lb/up<sup>@</sup> | RMSD |
| 4L9Q | teniposide | Pyrx Virtual Screening | Autodock Vina | -11,4 | 0 |  |
|  |  | Chimera | Autodock Vina | -12,1 | 0 | 3,64 Å |
|  |  | Emdock | AutoDock 4 | Binding affinity (kcal/mol)<br>-10,98 | Ki<br>8,95 nM | Ligand efficiency<br>-0,24 |
| Target | Drug | Docking programs | Docking software | Binding affinity (kcal/mol) | RMSD lb/up<sup>@</sup> | RMSD |
| 4LA0 | bicalutamide | Pyrx Virtual Screening | Autodock Vina | -9,9 | 0 |  |
|  |  | Chimera | Autodock Vina | -9,8 | 0 | 9,57 Å |
|  |  | Emdock | AutoDock 4 | Binding affinity (kcal/mol)<br>-4,94 | Ki<br>0,24 mM | Ligand efficiency<br>-0,17 |
| Target | Drug | Docking programs | Docking software | Binding affinity (kcal/mol) | RMSD lb/up<sup>@</sup> | RMSD |
| 4LB2 | idarubicin | Pyrx Virtual Screening | Autodock Vina | -9,4 | 0 |  |
|  |  | Chimera | Autodock Vina | -9,3 | 0 | 10,27 Å |
|  |  | Emdock | AutoDock 4 | Binding affinity (kcal/mol)<br>-8,04 | Ki<br>1,28 µM | Ligand efficiency<br>-0,22 |
| Target | Drug | Docking programs | Docking software | Binding affinity (kcal/mol) | RMSD lb/up<sup>@</sup> | RMSD |
| 4LB9 | etoposide | Pyrx Virtual Screening | Autodock Vina | -10 | 0 |  |
|  |  | Chimera | Autodock Vina | -9,4 | 0 |  |
|  |  | Emdock | AutoDock 4 | Binding affinity (kcal/mol)<br>-8,57 | Ki<br>522,58 nM | Ligand efficiency<br>-0,2 |

**Tab 3.**
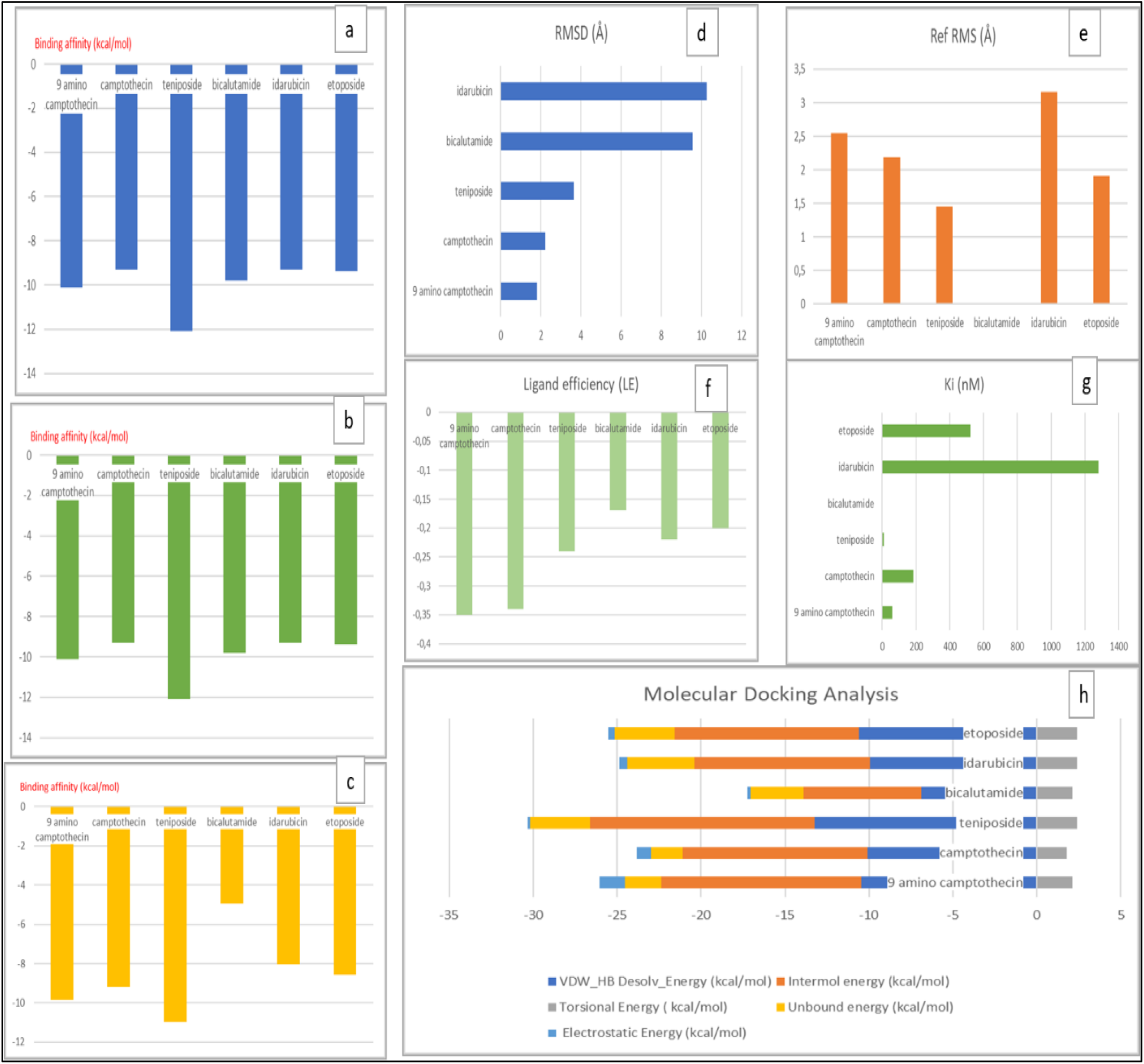
of Physical, Chemical and Biological Results of ID PDB: Crystal 4LA0 human serum albumin-Crystallised Bicalutamide b) ID PDB: Crystal 4L8U human serum albumin-Crystallised 9 amino Camptothecin c) ID PDB: Crystal 4L9K human serum albumin-Crystallised Camptothecin d) ID PDB: Crystal 4LB9 human serum albumin-Crystallised etoposide e) ID PDB: Crystal 4LB2 human serum albumin-Crystallised idarubicin f) ID PDB: Crystal 4L9Q human serum albumin-Crystallised teniposide

‐ **Binding Affinity** of 9-amino-camptothecin (ca.-10 kcal/mol), camptothecin (−9 kcal/mol) and teniposide (−11 kcal/mol)
‐ **RMSD Value** of 9 -amino-camptothecin (ca.1.8 Å), camptothecin (ca.2.2 Å) and teniposide (ca. 3.6 Å)
‐ **Ki Value**: 9 -amino-camptothecin (ca 59 nM), camptothecin (ca 183 nM) and teniposide (ca 9 nM)

#### Ligand efficiency

of 9 -amino-camptothecin(ca −0.35 kcal/mol), camptothecin (ca −0.34 kcal/mol) and teniposide (ca −0.24 kcal/mol)

##### Coordinates Docking for the center of the grid boxes

- ID PDB: 4L8U human serum albumin (HSA) - complexed with Crystallised 9 amino camptothecin >>Vina Search Space: center of the grid docking box grid docking box: center x=29.23; y=5.84; z=31.39; Dimensions (Angstroms) size x,y,z= 25/25/25 Å]
- ID PDB: 4L9K human serum albumin (HSA) - complexed with Crystallised camptothecin: >> Vina Search Space: center of the grid docking box: x=−9.85; y=−6.88; z=9.79; Dimensions (Angstroms) size x, y, z= 25/25/25 Å].
- ID PDB: 4L9Q human serum albumin (HSA) - complexed with Crystallised teniposide were >> Vina Search Space Vina Search Space : center of the grid docking box of the grid docking box: x=−13.107; y=−11.28; z=9.95; Dimensions (Angstroms) size x, y, z= 25/25/25 Å]
- ID PDB: 4LA0 human serum albumin (HSA) - complexed with Crystallised bicalutamide >> [central position and size of the grid docking box: x=−16.45; y=−12.02; z=8.18; Dimensions (Angstroms) size x, y, z= 25/25/25 Å].
- ID PDB: 4LB2 human serum albumin (HSA) - complexed with Crystallised idarubicin >> Vina Search Space: center of the grid docking box of the grid docking box: x=−14.40; y=−14.24; z=9.80; Dimensions (Angstroms) size x, y, z= 25/25/25 Å].
- ID PDB: 4LB9 human serum albumin (HSA) - complexed with Crystallised etoposide >> Search Space : center of the grid docking box: x=33.40; y=−2.79;z=16.69; Dimensions (Angstroms) size x,y,z= 25/25/25 Å].

In fig. 3 we explored the best three crystallized ligands in Human Serum Albumin. Indeed, 9 -amino Camptothecin, Camptothecin, and teniposide have excellent values in Affinity, RMSD, Ligand efficiency (LE), and Inhibitory constant Ki. Moreover, we observe a complete overlap, during the re-docking analysis phase, estimated by chimera Software. Therefore we have concluded that ID PDB Crystal 4L8U human serum albumin-Crystallised 9 -amino Camptothecin; ID PDB Crystal 4L9K human serum albumin-Crystallised Camptothecin and ID PDB Crystal 4L9Q human serum albumin-Crystallised teniposide be used as a possible referring protein to be compared with a target protein.

## 4. Conclusions

We concluded, through re-docking validation detailed analysis that only three of crystalized six oncology agents investageted in this work, in Human Serum Albumin protein, have excellent results. Indeed, 9 -amino Camptothecin, Camptothecin and teniposide in the Human Serum Albumin, have excellent values in <u>Affinity</u> (kcal/mol), Ligand efficiency (LE), and Inhibitory constant Ki. Moreover, we have observed a complete overlap, during the re-docking analysis phase. Therefore we believe that ID PDB Crystal 4L8U human serum albumin-Crystallised 9 -amino Camptothecin; ID PDB Crystal 4L9K human serum albumin-Crystallised Camptothecin and ID PDB Crystal 4L9Q human serum albumin-Crystallised teniposide can be taken as a reference protein compared against target proteins.

## Author contributions

I.V.F. conceived, designed and wrote the paper and performed the calculations and analyzed the data.

## Declaration of Competing Interest

The authors declare they have no potential conflicts of interest to disclose.

## Notes

### Competing Interest Statement

The authors have declared no competing interest.

## REFERENCES

1. Busher JT (1990). “Chapter 101: Serum Albumin and Globulin”. In Walker HK, Hall WD, Hurst JW (eds.). Clinical methods: the history, physical, and laboratory examinations (3rd ed.). Boston: Butterworths. ISBN 978-0409900774.

2. Komatsu T, Nakagawa A, Curry S, Tsuchida E, Murata K, Nakamura N, Ohno H (September 2009). “The role of an amino acid triad at the entrance of the heme pocket in human serum albumin for O(2) and CO binding to iron protoporphyrin IX”. Organic & Biomolecular Chemistry. 7 (18): 3836–41. doi:10.1039/b909794e3.

3. Curry S (August 2002). “Beyond expansion: structural studies on the transport roles of human serum albumin”. Vox Sanguinis. 83 Suppl 1: 315–9. doi:10.1111/j.1423-0410.2002.tb05326.x

4. Human Albumin structure in the Protein data bank; https://archive.is/20121128233355/160.114.99.91/astrojan/protein/pictures/albumin3.jpg

5. Sugio, A. Kashima, S. Mochizuki, M. Noda, K. Kobayashi, Crystal structure of human serum albumin at 2.5 Å resolution, Protein Engineering, Design and Selection, Volume 12, Issue 6, June 1999, Pages 439–446, https://doi.org/10.1093/protein/12.6.439

6. Carter, Daniel C., et al. “Three-dimensional structure of human serum albumin.” Science 244.4909 (1989): 1195–1198.

7. He, Xiao Min, and Daniel C. Carter. “Atomic structure and chemistry of human serum albumin.” Nature 358.6383 (1992): 209–215.

8. Kragh-Hansen, Ulrich. “Structure and ligand binding properties of human serum albumin.” Danish medical bulletin 37.1 (1990): 57–84.

9. Sudlow, G. D. J. B., D. J. Birkett, and D. N. Wade. “The characterization of two specific drug binding sites on human serum albumin.” Molecular Pharmacology 11.6 (1975): 824–832.

10. Fasano, Mauro, et al. “The extraordinary ligand binding properties of human serum albumin.” IUBMB life 57.12 (2005): 787–796.

11. Wang ZM, Ho JX, Ruble JR, Rose J, Rüker F, Ellenburg M, Murphy R, Click J, Soistman E, Wilkerson L, Carter DC. Structural studies of several clinically important oncology drugs in complex with human serum albumin. Biochim Biophys Acta. 2013 Dec;1830(12):5356–74. doi: 10.1016/j.bbagen.2013.06.032. Epub 2013 Jul 6. PMID: 23838380.

12. Chatterjee, Tanaya, et al. “Interaction of virstatin with human serum albumin: spectroscopic analysis and molecular modeling.” PloS one 7.5 (2012): e37468.

13. Kragh-Hansen, Ulrich. “Structure and ligand binding properties of human serum albumin.” Danish medical bulletin 37.1 (1990): 57–84.

14. KCarter, Daniel C., and Joseph X. Ho. “Structure of serum albumin.” Advances in protein chemistry. Vol. 45. Academic Press, 1994. 153–203.

15. Ghuman, Jamie, et al. “Structural basis of the drug-binding specificity of human serum albumin.” Journal of molecular biology 353.1 (2005): 38–52.

16. Madej T, Lanczycki CJ, Zhang D, Thiessen PA, Geer RC, Marchler-Bauer A, Bryant SH. “MMDB and VAST+: tracking structural similarities between macromolecular complexes. Nucleic Acids Res. 2014 Jan; 42(Database issue):D297–303

17. Wang ZM, Ho JX, Ruble JR, Rose J, Rüker F, Ellenburg M, Murphy R, Click J, Soistman E, Wilkerson L, Carter DC. Structural studies of several clinically important oncology drugs in complex with human serum albumin. Biochim Biophys Acta. 2013 Dec;1830(12):5356–74. doi: 10.1016/j.bbagen.2013.06.032. Epub 2013 Jul 6. PMID: 23838380.

18. National Center for Biotechnology Information (2020). PubChem Compound Summary for CID 2375, Bicalutamide. Retrieved December 27, 2020 from https://pubchem.ncbi.nlm.nih.gov/compound/Bicalutamide.

19. National Center for Biotechnology Information (2020). PubChem Compound Summary for CID 72402, 9-Aminocamptothecin. Retrieved December 27, 2020 from https://pubchem.ncbi.nlm.nih.gov/compound/9-Aminocamptothecin.

20. National Center for Biotechnology Information (2020). PubChem Compound Summary for CID 24360, Camptothecin. Retrieved December 27, 2020 from https://pubchem.ncbi.nlm.nih.gov/compound/Camptothecine.

21. National Center for Biotechnology Information (2020). PubChem Compound Summary for CID 36462, Etoposide. Retrieved December 27, 2020 from https://pubchem.ncbi.nlm.nih.gov/compound/Etoposide.

22. National Center for Biotechnology Information (2020). PubChem Compound Summary for CID 42890, Idarubicin. Retrieved December 27, 2020 from https://pubchem.ncbi.nlm.nih.gov/compound/Idarubicin.

23. National Center for Biotechnology Information (2020). PubChem Compound Summary for CID 452548, Teniposide. Retrieved December 27, 2020 from https://pubchem.ncbi.nlm.nih.gov/compound/Vumon.

24. Y. Xiang, L. Duan, Q. Ma, Z. Lv, Z. Ruohua, et al., Fluorescence spectroscopy and molecular simulation on the interaction of caffeic acid with human serum albumin, Luminescence (2016).

25. Mondal, Moumita, Krishna Ramadas, and Sakthivel Natarajan. “Molecular interaction of 2, 4-diacetylphloroglucinol (DAPG) with human serum albumin (HSA): the spectroscopic, calorimetric and computational investigation.” Spectrochimica Acta Part A: Molecular and Biomolecular Spectroscopy 183 (2017): 90–102

26. de Sousa, A.C.C., Combrinck, J.M., Maepa, K. et al. Virtual screening as a tool to discover new β-haematin inhibitors with activity against malaria parasites. Sci Rep 10, 3374 (2020). https://doi.org/10.1038/s41598-020-60221-0

27. Ma, Dik-Lung, Daniel Shiu-Hin Chan, and Chung-Hang Leung. “Molecular docking for virtual screening of natural product databases.” Chemical science 2.9 (2011): 1656–1665.

28. Ma, Dik-Lung, Daniel Shiu-Hin Chan, and Chung-Hang Leung. “Molecular docking for virtual screening of natural product databases.” Chemical science 2.9 (2011): 1656–1665.

29. Kellenberger, Esther, et al. “Comparative evaluation of eight docking tools for docking and virtual screening accuracy.” Proteins: Structure, Function, and Bioinformatics 57.2 (2004): 225–242.

30. Hawkins, Paul CD, A. Geoffrey Skillman, and Anthony Nicholls. “Comparison of shape-matching and docking as virtual screening tools.” Journal of medicinal chemistry 50.1 (2007): 74–82.

31. Pierri, Ciro Leonardo, Giovanni Parisi, and Vito Porcelli. “Computational approaches for protein function prediction: a combined strategy from multiple sequence alignment to molecular docking-based virtual screening.” Biochimica et Biophysica Acta (BBA)-Proteins and Proteomics 1804.9 (2010): 1695-1712.

32. Meloun B, Morávek L, Kostka V. Complete amino acid sequence of human serum albumin. FEBS Lett. 1975 Oct 15;58(1):134–7. doi: 10.1016/0014-5793(75)80242-0. PMID: 1225573.

33. Valdés-Tresanco, M.S., Valdés-Tresanco, M.E., Valiente, P.A. et al. AMDock: a versatile graphical tool for assisting molecular docking with Autodock Vina and Autodock4. Biol Direct 15, 12 (2020). https://doi.org/10.1186/s13062-020-00267-2

34. Abadzapatero, C; Metz, J (2005). “Ligand efficiency indices as guideposts for drug discovery”. Drug Discovery Today. 10 (7): 464–469. doi:10.1016/S1359-6446(05)03386-6

35. https://pyrx.sourceforge.io/

36. Small-Molecule Library Screening by Docking with PyRx. Dallakyan S, Olson AJ. Methods Mol Biol. 2015;1263:243-50. The full-text is available at https://www.researchgate.net/publication/273954875_SmallMolecule_Library_Screening_by_Docking_with_PyRx

37. http://autodock.scripps.edu/resources/references

38. Morris, G. M., Huey, R., Lindstrom, W., Sanner, M. F., Belew, R. K., Goodsell, D. S. and Olson, J. (2009) Autodock4 and AutoDockTools4: automated docking with selective receptor flexiblity. J. Computational Chemistry 2009, 16: 2785–91.

39. Morris, G. M., Goodsell, D. S., Halliday, R.S., Huey, R., Hart, W. E., Belew, R. K. and Olson, A. J. (1998), Automated Docking Using a Lamarckian Genetic Algorithm and and Empirical Binding Free Energy Function J. Computational Chemistry, 19: 1639–1662.

40. Ruth Huey., Morris, G. M., Olson, A. J. and Goodsell, D. S. (2007), A Semiempirical Free Energy Force Field with Charge-Based Desolvation J. Computational Chemistry, 28: 1145–1152.

41. Morris, G. M., Goodsell, D. S., Huey, R. and Olson, A. J. (1996), Distributed automated docking of flexible ligands to proteins: Parallel applications of AutoDock 2.4 J. Computer-Aided Molecular Design, 10: 293–304.

42. Goodsell, D. S., Morris, G. M. and Olson, A. J. (1996), Automated Docking of Flexible Ligands: Applications of AutoDock J. Mol. Recognition, 9: 1–5.

43. O. Trott, A. J. Olson, AutoDock Vina: improving the speed and accuracy of docking with a new scoring function, efficient optimization and multithreading, Journal of Computational Chemistry 31 (2010) 455–461

44. https://discover.3ds.com/discovery-studio-visualizer-download

45. BIOVIA, Dassault Systèmes, [Software product name], [Software version], San Diego: Dassault Systèmes, [Year]. Please note that all current BIOVIA software products start with the company name, and should be referenced as such. All products can be found at https://3ds.com/products-services/biovia/products

46. https://www.cgl.ucsf.edu/chimera/

47. Pettersen, E.F., Goddard, T.D., Huang, C.C., Couch, G.S., Greenblatt, D.M., Meng, E.C., and Ferrin, T.E. “UCSF Chimera - A Visualization System for Exploratory Research and Analysis.” J. Comput. Chem. 25(13):1605–1612 (2004).

48. Valdés-Tresanco, M.S., Valdés-Tresanco, M.E., Valiente, P.A. et al.. AMDock: a versatile graphical tool for assisting molecular docking with Autodock Vina and Autodock4. Biol Direct 15, 12 (2020). https://doi.org/10.1186/s13062-020-00267-2

49. Pantsar, T.; Poso, A. Binding Affinity via Docking: Fact and Fiction. Molecules 2018, 23, 1899. Protein–Ligand Docking - Profacgen

50. Lengauer T, Rarey M (Jun 1996). “Computational methods for biomolecular docking”. Current Opinion in Structural Biology. 6 (3): 402–6. doi:10.1016/S0959-440X(96)80061-3

51. Kitchen DB, Decornez H, Furr JR, Bajorath J (Nov 2004). “Docking and scoring in virtual screening for drug discovery: methods and applications”. Nature Reviews. Drug Discovery. 3 (11): 935–49. doi:10.1038/nrd1549

52. Henrich, S., Feierberg, I., Wang, T., Blomberg, N. and Wade, R.C. (2010), Comparative binding energy analysis for binding affinity and target selectivity prediction. Proteins, 78: 135–153. https://doi.org/10.1002/prot.22579

46. Huey, Ruth, Garrett M. Morris, Arthur J. Olson, and David S. Goodsell. “A Semiempirical Free Energy Force Field with Charge-Based Desolvation.” Journal of Computational Chemistry 28, no. 6 (April 30, 2007): 1145–52. doi:10.1002/jcc.20634.

53. Molecular docking, estimating free energies of binding, and AutoDock’s semi-empirical force field”. Sebastian Raschka’s Website. 2014-06-26. Retrieved 2016-06-07.

54. Coutsias EA, Seok C, Dill KA (2004). “Using quaternions to calculate RMSD”. J Comput Chem. 25 (15): 1849–1857. doi:10.1002/jcc.20110

